# A *Pax3* lineage gives rise to transient haematopoietic progenitors

**DOI:** 10.1101/2024.06.26.600366

**Authors:** Giovanni Canu, Rosamaria Correra, Guillermo Díez Pinel, Raphaël F. P. Castellan, Laura Denti, Alessandro Fantin, Christiana Ruhrberg

**Author notes:** Corresponding author: Professor Christiana Ruhrberg, UCL Institute of Ophthalmology, University College London, 11-43 Bath Street, London EC1V 9EL, United Kingdom.

## Abstract

During embryonic development, muscle tissues, skin, and a subset of vascular endothelial cells arise from *Pax3-*expressing embryonic progenitors defined as paraxial mesoderm. By contrast, haemogenic potential is well established for extra-embryonic mesoderm and intra-embryonic lateral plate mesoderm which do not express *Pax3*. To date, it is not known whether the haematopoietic system also contains *Pax3* lineage cells. Here, we show that the mouse foetal liver and foetal circulation contain a transient population of *Pax3* lineage cells with hallmarks of haematopoietic progenitors and the potential to generate both myeloid and erythroid cells. We propose that *Pax3* lineage haematopoietic cells should be investigated to better understand normal haematopoietic development from different mesodermal derivatives. Further, genetic alterations of *Pax3* lineage haematopoietic cells should be investigated for their potential to cause haematopoietic malignancies.

## INTRODUCTION

Deducing cell lineages in developmental biology research is commonly achieved with genetic lineage tracing in the mouse via Cre/LoxP-mediated DNA recombination^1^, whereby Cre recombinase expressed from a cell type-specific mouse promoter irreversibly excises a stop codon upstream of a fluorescent reporter^2^, such as *Rosa*^*Egfp*^, *Rosa*^*Yfp*^ or *Rosa*^*tdTom*^. Using this method with the *Pax3*^*Cre*^ knock-in allele^3^, it has been shown that muscle cells and a subset of endothelial cells arise from *Pax3-*expressing paraxial mesoderm, whereas melanocytes and a subset of neurons arise from *Pax3*-expressing neural crest^4–8^. Recently, bulk RNA sequencing (RNA-seq) of *Pax3*^*Cre*^ lineage-traced embryonic mouse limbs identified unexplained transcripts for ‘immune system-related genes’ alongside the expected musculoskeletal and neuronal markers typical of paraxial mesoderm and neural crest derivatives^9^.

During embryonic development, haematopoiesis occurs in temporally and spatially overlapping waves that originate from well-defined tissues^10^. Extra-embryonic mesoderm gives rise to yolk sac haemogenic endothelium that transitions into pro-definitive haematopoietic progenitors, including erythro-myeloid progenitors (EMPs) and lympho-myeloid progenitors (LMPs)^10–12^. By contrast, intra-embryonic lateral plate mesoderm produces haemogenic endothelium in the dorsal aorta that transitions into definitive haematopoietic stem cells (HSCs)^10,13,14^. Neither type of mesoderm with haematopoietic potential is known to express *Pax3*, which instead is considered to be selectively expressed in paraxial mesoderm and neural crest^4–8^.

Here, we show that the mouse foetal liver and foetal circulation contain a transient population of *Pax3* lineage cells with hallmarks of haematopoietic progenitors and the potential to generate both myeloid and erythroid cells. Our findings raise the possibility that haemogenic potential, observed for extra-embryonic mesoderm and intra-embryonic lateral plate mesoderm, may also extend to paraxial mesoderm.

## RESULTS AND DISCUSSION

Re-analysis of bulk RNA-seq data^9^ from FACS-isolated embryonic day (E) 12.5 mouse limbs genetically lineage traced with *Pax3*^*Cre*^*;Rosa*^*Egfp*^ detected the expected transcripts for the paraxial mesoderm-derived musculoskeletal (*e*.*g*., *Myf5, Myog*) and endothelial (*e*.*g*., *Cdh5, Kdr*) cell lineages, but also transcripts typical of haematopoietic progenitors (*e*.*g*., *Kit, Vav1, Runx1*), myeloid cells (*e*.*g*., *Csf1r, Cd68*) and erythroid cells (*e*.*g*., *Hbb-bh1, Hbb-bs*; **Fig. 1A**). Transcription factors required for lymphoid differentiation were also detected (*e*.*g*., *Ebf1, Tcf7*), but not markers of mature lymphocytes (*e*.*g*., *Cd3, Cd4, Cd19*; **Fig. 1A**). The finding suggests that a *Pax3* lineage may give rise to a subset of haematopoietic cells.

**Fig. 1.**
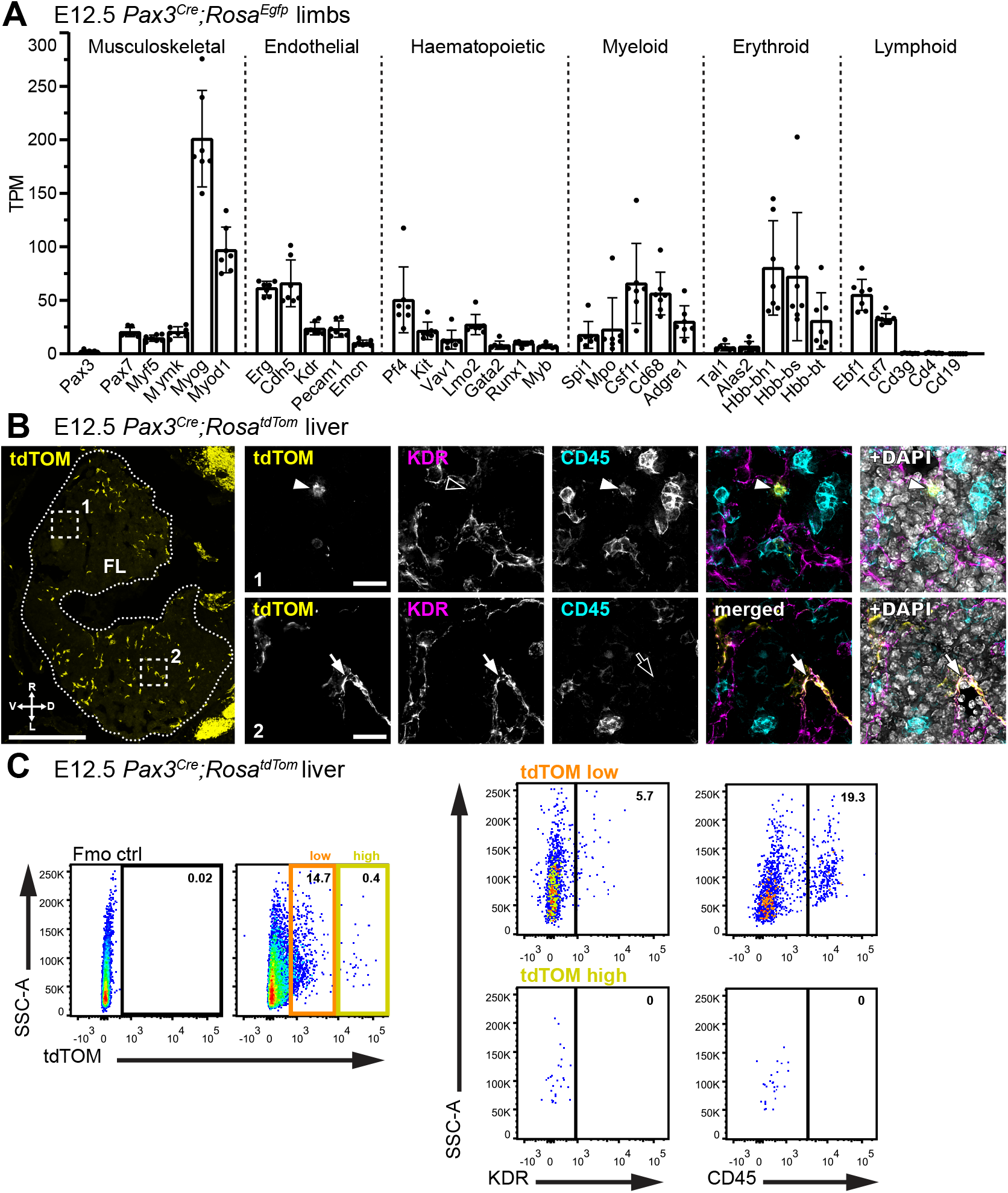
*Pax3* lineage endothelial and haematopoietic cells in the foetal liver and blood. **A** Bulk RNA-seq analysis of EGFP+ cells from E12.5 *Pax3*^*Cre*^*;Rosa*^*Egfp*^ mouse limbs (ENA project PRJNA422253) for transcripts from the indicated genes; n = 7 embryos. **B** Representative immunofluorescence staining with the indicated markers of a E12.5 *Pax3*^*Cre*^*;Rosa*^*tdTom*^ embryo section at the level of the foetal liver (FL, outlined with dots); n = 3 embryos. Scale bar: 500 µm. The dorsal (D), ventral (V), right (R) and left (L) side of the embryo are indicated. The dashed squares indicate two areas shown at higher magnification in the adjacent panels to visualise tdTOM+CD45+ haematopoietic cells (arrowheads in panel 1) and tdTOM+KDR+ endothelial cells (arrows in panel 2). Scale bar: 20 µm. **C** Flow cytometry analysis including representative dot plots of E12.5 *Pax3*^*Cre*^*;Rosa*^*tdTom*^ liver, showing the proportion of endothelial and haematopoietic cells in the tdTOM low and tdTOM high fractions; n = 5 embryos for KDR, n = 10 embryos for CD45.

We next examined the *Pax3* lineage trace in the mouse foetal liver, which harbours haematopoietic progenitors from midgestation until birth to support foetal haematopoiesis^10^. Immunofluorescence staining identified *Pax3* lineage KDR+ endothelial and CD45+ haematopoietic cells in E12.5 *Pax3*^*Cre*^*;Rosa*^*tdTom*^ liver (**Fig. 1B**). Flow cytometry corroborated the presence of *Pax3* lineage KDR+ and CD45+ cells in E12.5 liver (**Fig. 1C**; 46.98 ± 11.36% of KDR+ endothelial cells and 25.70 ± 7.15% of CD45+ haematopoietic cells in the E12.5 liver were lineage-traced with *Pax3*^*Cre*^). *Pax3* lineage cells were also identified in the circulation but were less abundant than in the liver (E12.5 liver: 15.83 ± 5.09%; E12.5 blood: 2.99 ± 3.20%; **Fig. 2A,B**). In both the liver and blood, the proportion of *Pax3* lineage cells peaked at E12.5 and then rapidly declined (**Fig. 2A,B**). At E12.5, 16.89 ± 3.46% of *Pax3* lineage cells in the liver and 9.64 ± 4.88% in the blood were CD45+ haematopoietic cells (**Fig. 2C,D**). The *Pax3* lineage-traced populations at both sites also included TER119+ erythroid cells (E12.5 liver: 39.31 ± 5.73%; E12.5 blood: 31.84 ± 24.18%; **Fig. 2C,D**;). Moreover, a subset of CD45+ cells co-expressed the haematopoietic progenitor marker KIT (**Fig. 2C,D**). Consistent with haematopoietic progenitors being enriched in the foetal liver at midgestation^10^, *Pax3* lineage CD45+KIT+ cells were more abundant in the liver than blood (E12.5 liver: 12.27 ± 2.00%; E12.5 blood: 3.06 ± 2.73%; **Fig. 2C,D**).

**Fig. 2.**
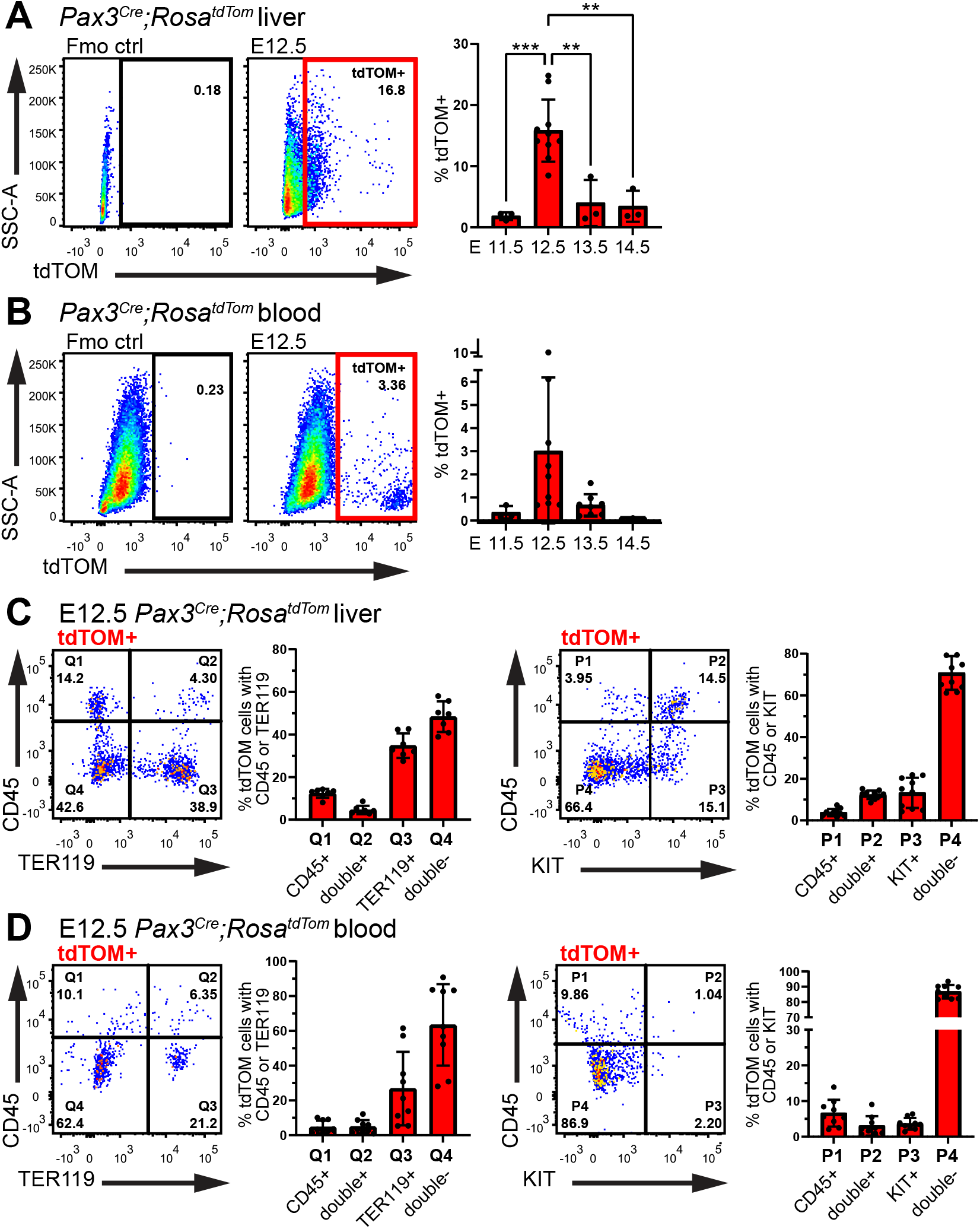
Transience of *Pax3* lineage haematopoietic cells in the foetal liver and blood. Flow cytometry analysis of *Pax3*^*Cre*^*;Rosa*^*tdTom*^ liver (**A**,**C**) and blood (**B**,**D**) including representative dot plots of E12.5 tdTOM+ cells with quantitative analysis of E11.5-14.5 tdTOM+ cells (**A**,**B**), and quantitative analysis of E12.5 tdTOM+ CD45+ TER119+ KIT+ cells (**C**,**D**). Data are shown as mean ± standard deviation (SD); each data point represents the value from an individual embryo; liver: n = 3 E11.5, n = 10 E12.5, n = 3 E13.5, n= 3 E14.5; blood: n = 3 E11.5, n = 9 E12.5, n = 8 E13.5, n= 3 E14.5. *P<0.5, **P<0.01, ***P<0.001, ****P<0.0001 (one-way ANOVA).

Next, we quantified the contribution of *Pax3* lineage cells to the developing haematopoietic system at E12.5. In the liver, more than 25% of CD45+ cells overall and CD45+KIT+ progenitors were *Pax3* lineage-traced (CD45+: 25.70 ± 7.15%; CD45+KIT+: 26.49 ± 7.47%; **Fig. 3A**). The proportion of *Pax3* lineage-traced CD45+KIT+ cells in the blood was approximately half of that in the liver at E12.5 (12.63 ± 6.89%, **Fig. 3B**), again consistent with haematopoietic progenitor enrichment in the midgestation foetal liver^10^. *Pax3* lineage TER119+ erythroid cells were present in the E12.5 liver but rare in the blood (liver: 10.71 ± 2.38%; blood: 0.52 ± 0.27%; **Fig. 3A,B**). Considering that most TER119+ cells in the E12.5 circulation are primitive erythrocytes^10^, foetal liver presence of *Pax3* lineage TER119+ cells, but their absence from the circulation, suggests that *Pax3* lineage haematopoietic cells are not derived from the so-called primitive haematopoietic wave. Instead, these findings suggest that *Pax3* lineage TER119+ cells arise locally in the foetal liver, as described for erythroid cell production from pro-definitive EMPs at midgestation^10^.

**Fig. 3.**
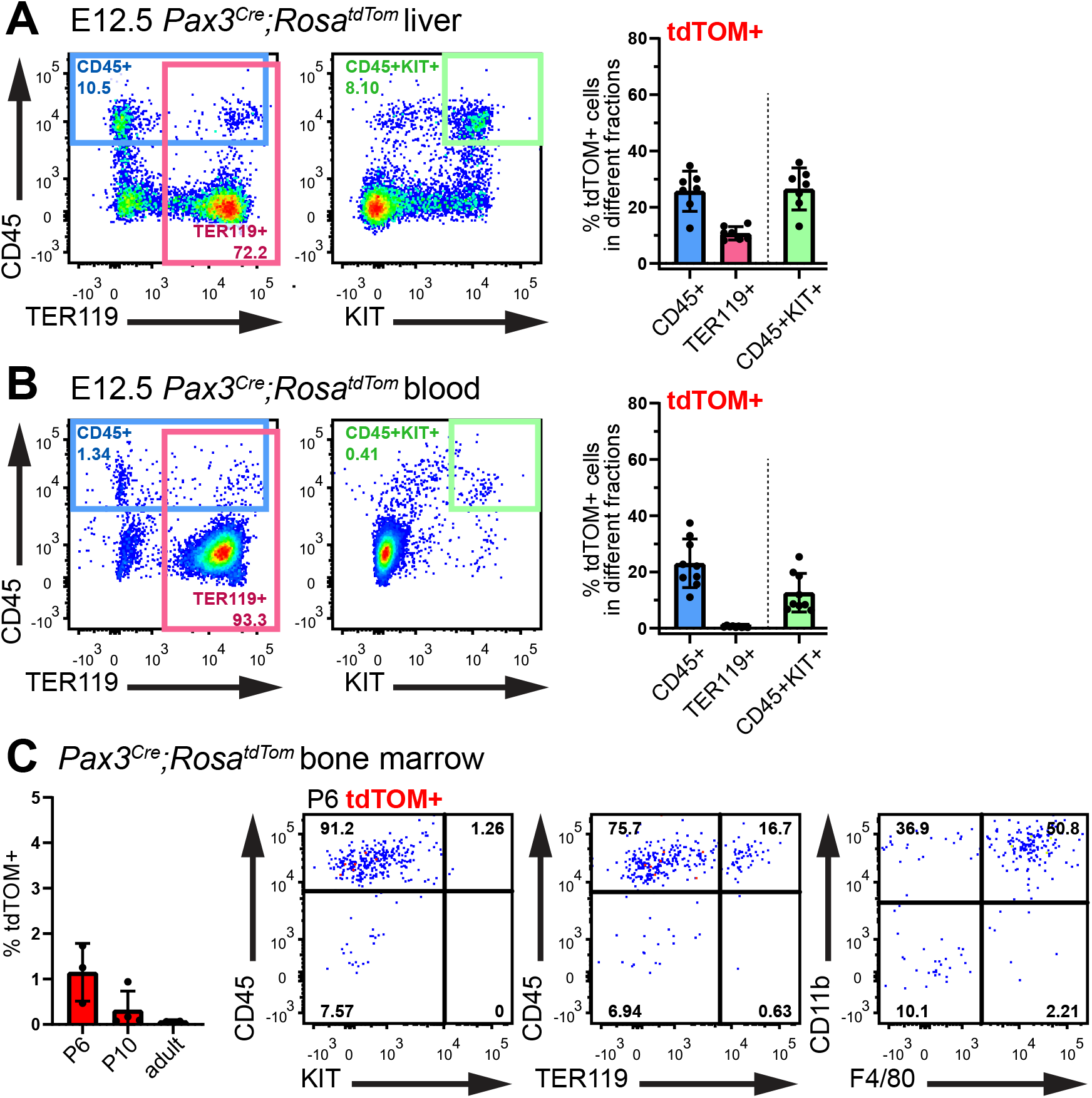
The *Pax3* lineage contributes haematopoietic progenitors to the foetal liver and blood but not post-natal bone marrow. **A**,**B** Flow cytometry analysis of E12.5 *Pax3*^*Cre*^*;Rosa*^*tdTom*^ liver (**A**) and blood (**B**), including representative dot plots and quantitative analysis of the fractions of all CD45+, TER119+ and CD45+KIT+ cells expressing tdTOM. Data are shown as mean ± SD; each data point represents the value from an individual embryo; liver: n = 7 E12.5; blood: n = 9. **C** Flow cytometry analysis of *Pax3*^*Cre*^*;Rosa*^*tdTom*^ bone marrow, including quantitative analysis of P6, P10 and adult (10 months old) tdTOM+ cells, and representative dot plots of P6 tdTOM+ cells. Data are shown as mean ± SD; P6: n = 3, P10: n = 4; adult: n = 2.

During foetal development, HSCs home to the foetal liver from around E12.5 onwards, before they seed the bone marrow to sustain adult haematopoiesis^10^. Therefore, we investigated whether *Pax3* lineage haematopoietic progenitors also migrated to the bone marrow. Flow cytometry of postnatal day 6 (P6) *Pax3*^*Cre*^*;Rosa*^*tdTom*^ bone marrow detected rare *Pax3* lineage cells (1.15 ± 0.64%) that were mostly CD45+CD11b+F4/80+ macrophages and CD45+CD11b+F4/80-myeloid cells; by contrast, *Pax3* lineage KIT+CD45+ progenitors or TER119+CD45-erythrocytes were not observed (**Fig. 3C**). Fewer *Pax3* lineage cells were detected in P10 than P6 bone marrow, and they were undetectable in adult bone marrow (**Fig. 3C**). Absent adult bone marrow colonisation is inconsistent with *Pax3* lineage haematopoietic progenitors being a subset of HSCs but is consistent with a pro-definitive, EMP-like identity.

To determine the haematopoietic potential of *Pax3* lineage progenitors, we isolated E12.5 liver CD45+KIT+ cells lacking differentiated lineage markers for colony-forming unit (CFU) assays, in which they gave rise to multilineage CFU-GEMM along with erythroid and myeloid colonies (**Fig. 4A-C**). Detecting uniform tdTOM expression in differentiated CFU colonies excluded that the haematopoietic potential of explanted progenitors was derived from contaminating tdTOM negative cells (**Fig. 4D**). Further, the *Pax3* lineage CFU colonies expressed blood lineage markers consistent with differentiated erythroid and myeloid cells (**Fig. 4E**). Finding that *Pax3* lineage haematopoietic progenitors have erythroid and myeloid differentiation potential, in the context of their transient presence in the foetal liver but absence from bone marrow, again agrees with the idea that these cells are more similar to pro-definitive EMPs than definitive HSCs. The residual presence of *Pax3* lineage macrophages in the neonatal bone marrow, in the absence of *Pax3* lineage haematopoietic progenitors, is reminiscent of EMP-derived macrophages, which persist as tissue-resident in adults in an organ-specific manner even though their progenitors are only transient^10^. Nevertheless, future studies should investigate whether *Pax3* lineage macrophages also persist in adult organs.

**Fig. 4.**
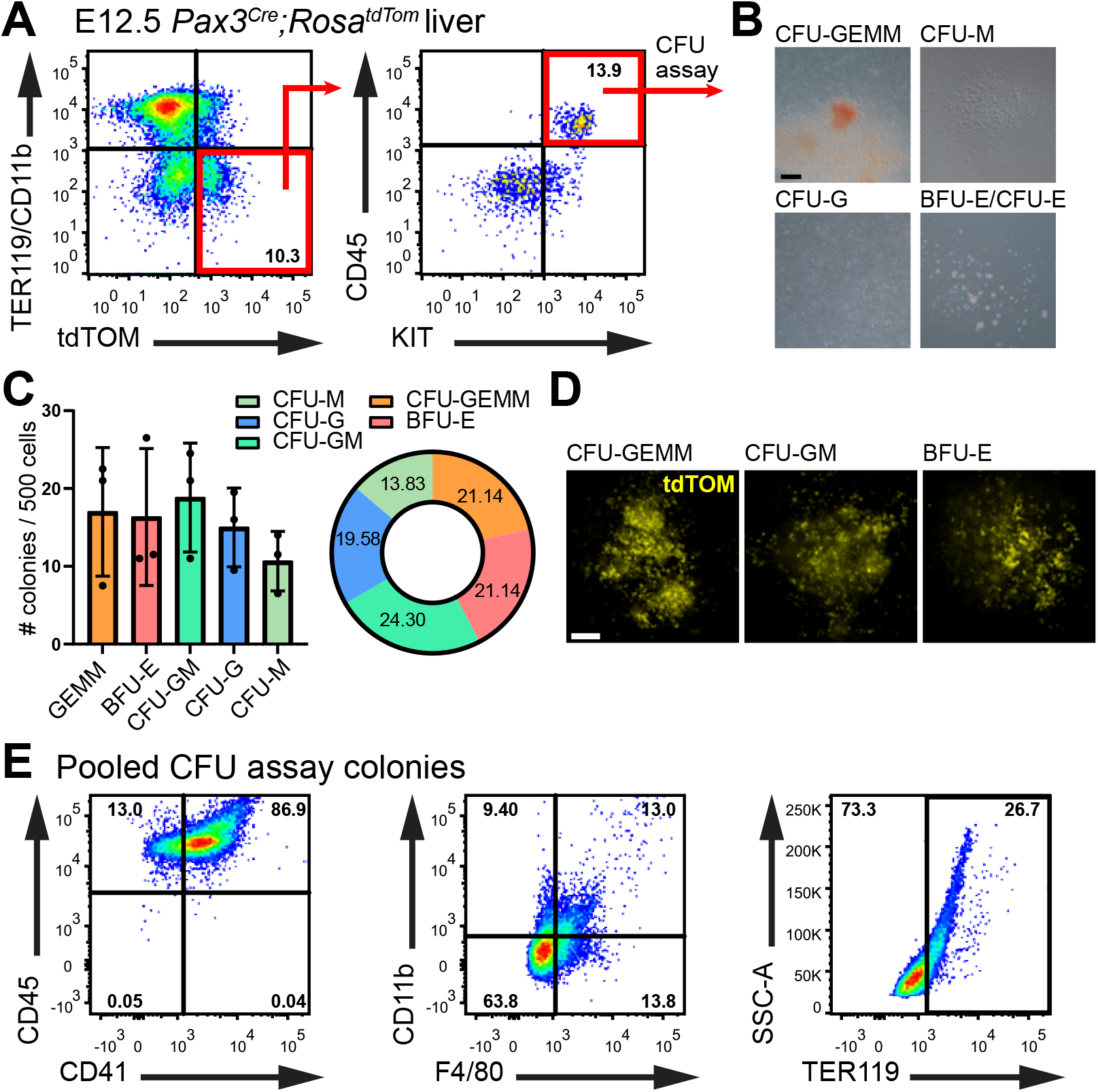
*Pax3* lineage cells have hallmarks of multipotent haematopoietic progenitors. *Pax3*-lineage (tdTOM+) TER119- CD11b- KIT+ CD45+ haematopoietic progenitors were isolated from E12.5 *Pax3*^*Cre*^*;Rosa*^*tdTom*^ liver for CFU assays. **A** Representative dot plots highlight sorted cells (red squares). **B** Representative brightfield images of CFU assay colonies; CFU-GEMM, granulocyte-erythrocyte-macrophage-megakaryocyte; BFU-E, burst-forming unit erythrocyte; CFU-GM, granulocyte-macrophage; CFU-G, granulocyte; CFU-M, macrophage. Scale bar: 200 µm. **C** Quantitative analysis of colonies scored after 12 days; bar plot data are shown as mean ± SD; each data point represents a pooled litter of 5-7 embryos (n = 3 litters); pie chart shows mean values as percentage of total colonies obtained. **D** Fluorescence image of a representative CFU colonies. **E** Representative dot plots of flow cytometry analysis from pooled CFU colonies with the indicated markers (n = 3).

As *Pax3* is an established marker of paraxial mesoderm and neural crest, both populations might be considered as potential sources of *Pax3* lineage haematopoietic progenitors. In particular, it is conceivable that these progenitors might arise from a subset of paraxial mesoderm-derived endothelial cells with haemogenic potential, in a process analogous to that described for extra-embryonic mesoderm-derived haemogenic endothelium in the yolk sac or lateral plate mesoderm-derived haemogenic endothelium in the dorsal aorta^11,14,15^. Alternatively, *Pax3* may be expressed in a distinct and hitherto unrecognised source of progenitors with haematopoietic potential that may or may not pass through an endothelial intermediate, in which case *Pax3*-mediated lineage tracing would be insufficient to attribute cellular origins to paraxial mesoderm.

We have not investigated whether *Pax3* lineage haematopoietic progenitors may be involved in haematopoietic malignancies. Nevertheless, it is conceivable that genetic alterations that cause leukemic transformation in haematopoietic stem and progenitor cells might also affect *Pax3* lineage haematopoietic progenitors. Furthermore, the *Pax3* lineage origin of these transient progenitors raise the possibility that they may also be affected by genetic alterations that are not commonly taken into consideration for haematological studies. In this context, we note that a t(2;13)(q35;q14) genetic translocation produces a tumorigenic PAX3-FOXO1 fusion protein in patients with paediatric alveolar rhabdomyosarcoma (aRMS)^16,17^. This mutation increases the proliferation of *Pax3*-expressing cells whilst inhibiting their terminal differentiation^18^ and also promotes the trans-differentiation of endothelial cells towards a myogenic fate^19^. Thus, it should be examined whether PAX3-FOXO1 could also induce myogenic trans-differentiation in haematopoietic cells, or whether it might alter haematopoietic development via an endothelial-derived *Pax3* lineage haematopoietic progenitor. These are interesting considerations, because some aRMS patients with PAX3-FOXO1 fusion protein have a bone marrow phenotype resembling acute leukaemia^20^, which is currently thought to result from metastatic aRMS infiltrating the bone marrow, even in cases with no identifiable primary tumor^21^. Existing mouse models^22,23^ could be used to determine whether such translocation can cause primary haematopoietic malignancies.

### Conclusion

Here, we have identified novel *Pax3* lineage cells with hallmarks of transient embryonic haematopoietic progenitors. Our results should open novel lines of investigations to determine the origins and role of *Pax3* lineage progenitors during normal haematopoietic development, and their relevance for haematological malignancies.

## METHODS

### Transcriptomic studies

Raw bulk RNA-seq reads from *Pax3*^*Cre*^*;Rosa*^*Egfp*^ mouse embryo limbs (PRJNA422253)^9^ were downloaded from the European Nucleotide Archive, aligned to GRCm39 and annotated using Mus_musculus.GRCm39.110.gtf at http://www.ensembl.org/info/data/ftp/index.html. Transcripts per million (TPM) values were plotted in Prism 9 (GraphPad).

### Animal procedures and tissue staining

Animal procedures were performed according to Animal Welfare Ethical Review Body (AWERB) and UK Home Office guidelines. C57BL/6J mice carrying the *Pax3*^*Cre*^ knock-in allelle^3^ were timed-mated to mice carrying a *Rosa*^*tdTom*^ recombination reporter^2^. Cryosections of formaldehyde-fixed E12.5 liver were blocked in PBS containing 10% serum-free protein block (DAKO), 2.5% BSA, and 0.1% Triton X-100 before staining with antibodies for KDR (R&D Systems #AF644), CD45 (BD Biosciences #550539), and RFP (MBL #PM005) followed by donkey Fab fragments (Stratech Jackson): Alexa Fluor 647-conjugated anti-goat (#705-606-147), Alexa Fluor 488-conjugated anti-rat (#712-547-003) and Cy3-conjugated anti-rabbit (#711-166-152). DAPI-counterstained sections were imaged on a Ti2 microscope with NIS-Elements automatic deconvolution (Nikon).

### Flow cytometry

Embryos were collected in ice-cold FACS buffer: RPMI with 2.5% foetal bovine serum (ThermoFisher), 100 µg/ml heparin and 50 µg/ml DNAse I (Sigma). Extraembryonic and head tissues were removed for blood collection before livers were dissected and dissociated in FACS buffer with 100 µg/ml collagenase/dispase (Sigma) for 20 min and TrypLE (Gibco) for 3 min. Single cell suspensions from foetal liver, foetal blood or post-natal femur and tibia bone marrow, were incubated with Fc block (Biolegend) for 30 min before labelling with the following antibodies (Biolegend): KIT (clone 2B8), CD45 (clone 30-F11), TER119 (clone TER-119), CD11b (clone M1/70), CD41 (clone MWReg30), F4/80 (clone BM8). Live cells were analysed using a Fortessa X-20 (BD Biosciences) or sorted on a FACSAriaIII (BD Biosciences). Data were analysed using FlowJo VX (FlowJo) and Prism 9 (GraphPad).

### Haematopoietic differentiation assay

Haematopoietic differentiation was tested in colony-forming unit assays using Methocult GF-M3434 (STEMCELL Technologies), following manufacturer instructions.

## ACKNOWLEDGEMENTS

We thank the staff of the Biological Resources, FACS and Imaging Facilities at the UCL Institute of Ophthalmology. This research was supported by grants from the Wellcome [205099/Z/16/Z] to CR, the British Heart Foundation [PG/18/85/34127 and PG/23/11301] to CR and [FS/19/14/34170] to RC and the Fondazione Associazione Italiana per la Ricerca sul Cancro (AIRC) [22905] to AF.

## AUTHORSHIP CONTRIBUTIONS

GC, RC, AF and CR conceived and designed the study. GC and CR co-wrote the manuscript. GC and LD performed genetic crosses and genotyping. GC, RC and RFPC performed experiments and analysed data. GDP and RFPC performed bioinformatic analyses. CR supervised the project. All authors read and approved the submitted manuscript.

